# Canonical Wnt Signaling is Involved in Anterior Regeneration of the Annelid *Aeolosoma viride*

**DOI:** 10.1101/2020.03.01.972448

**Authors:** Cheng-Yi Chen, Wei-Ting Yueh, Jiun-Hong Chen

## Abstract

Annelids are regenerative animals, but the underlying mechanisms await to be discovered. Because Wnt pathway is involved in animal regeneration to varying extents, we used *Aeolosoma viride* to interrogate whether and how this pathway plays a role in annelid anterior regeneration. We found that the expression of *wnt4*, *β-catenin* and nuclear-localized β-catenin protein were up-regulated during blastemal formation and down-regulated as anterior structures gradually reformed. Consistent with potential Wnt activities in the blastema, treatments with either Wnt pathway activator (azakenpaullone) or inhibitor (XAV939) inhibited head regeneration, which further supports a role of Wnt pathway during anterior regeneration. Detailed tissue-level examines demonstrated that wound closure and blastemal cell proliferation were impaired by over-activating the pathway, and that neuronal and musculature differentiation were affected under Wnt inhibition. Combined, gene expression and chemical inhibitor data suggest the presence of dynamic Wnt activities at different anterior regeneration stages: an initial low activity may be required for wound closure, and the following activation may signal blastemal formation and cell differentiation. In a nutshell, we propose that the canonical Wnt signaling regulates blastemal cellular responses during annelid regeneration.

## Introduction

Among regeneration models, researches from hydra and planarian have uncovered many cellular and molecular mechanisms (King & Newmark, 2012; E. M. Tanaka & Reddien, 2011). To gain more perspectives into animal regeneration and to relate to broader evolutionary implications, researches in other emerging models are indispensable. Annelids are known for regeneration capabilities and some underlying mechanisms are evolutionarily shared with other animals. For example, similar to the neoblasts of planarian (King & Newmark, 2012; E. M. Tanaka & Reddien, 2011), *Enchytraeus japonensis* and *Pristina leidyi* maintain stem cells in every segment for regeneration (Myohara, 2012; Yoshida-Noro & Tochinai, 2010; Zattara, Turlington, & Bely, 2016). Proliferative stem cells of planarian and *Eisenia foetida* express pluripotent factors, such as *oct4*, *nanog* and *sox2* (Onal et al., 2012; Zheng, Shao, Diao, Li, & Han, 2016). Consistent with the requirement of innervation for vertebrate limb regeneration (Bryant, Endo, & Gardiner, 2002), diverted neuron fiber induces ectopic axis in *Eisenia* annelids (Bely, 2014). However, whether any molecular mechanism regulates annelid regeneration is not clear yet (Bely, 2014).

Canonical Wnt signaling pathway (i.e. the Wnt/β-catenin pathway) plays major roles in various regenerative models. In hydra, injury induced apoptosis triggers the secretion of wnt3 from dying cells and activates the oral pole regeneration (Chera et al., 2009). Knocking-down Wnt pathway genes affect the anterior-posterior axial registration during planarian regeneration (Augustin et al., 2006; Broun, Gee, Reinhardt, & Bode, 2005; Gurley, Rink, & Alvarado, 2008; Iglesias, Gomez-Skarmeta, Saló, & Adell, 2008; King & Newmark, 2012; Lengfeld et al., 2009; Petersen & Reddien, 2008; E. Tanaka & Weidinger, 2008). Additionally, activating Wnt signaling enhances fin and limb regeneration in zebrafish and axolotl (Kawakami et al., 2006; Yokoyama, Ogino, Stoick-Cooper, Grainger, & Moon, 2007). In *Xenopus laevis,* activation of Wnt pathway rejuvenates the regenerative ability of aged limbs (Yokoyama et al., 2007). Therefore, Wnt pathway is likely to regulate regeneration in other models.

Here, we used an emerging annelid regeneration model *Aeolosoma viride* to interrogate the involvement of canonical Wnt pathway in annelid regeneration (C.-P. Chen et al., 2020). *A. viride* belongs to Aeolosomaitidae, Annelida, and is phylogenetically classified as “clitellate-like polychaetes” by morphological or molecular evidences (Hessling & Purschke, 2000; Zrzavý, Říha, Piálek, & Janouškovec, 2009). *A. viride* is a freshwater annelid, typically ranges 2-3 mm in length and 0.1-0.2 mm in width with brown spots scatter on its semitransparent epithelium. In lab conditions, *A. viride* only reproduce asexually by adding posterior segments, and the progeny detaches from the parent worm via paratomic fission (Falconi, Gugnali, & Zaccanti, 2015). Taking the advantage of its semitransparent appearance, we are able to directly observe its internal organs, such as horseshoe-shaped mouth and the digestive tract (Fig. 1A). In our preliminary tests, we found *A. viride* can robustly regenerate anterior segments in 5 days (C.-P. Chen et al., 2020). The posterior-most pygidium segment is hardly distinguishable from posterior wounds, resulting in biased assessment to successful regeneration. Therefore, we focused on anterior regeneration here.

**Fig. 1.**
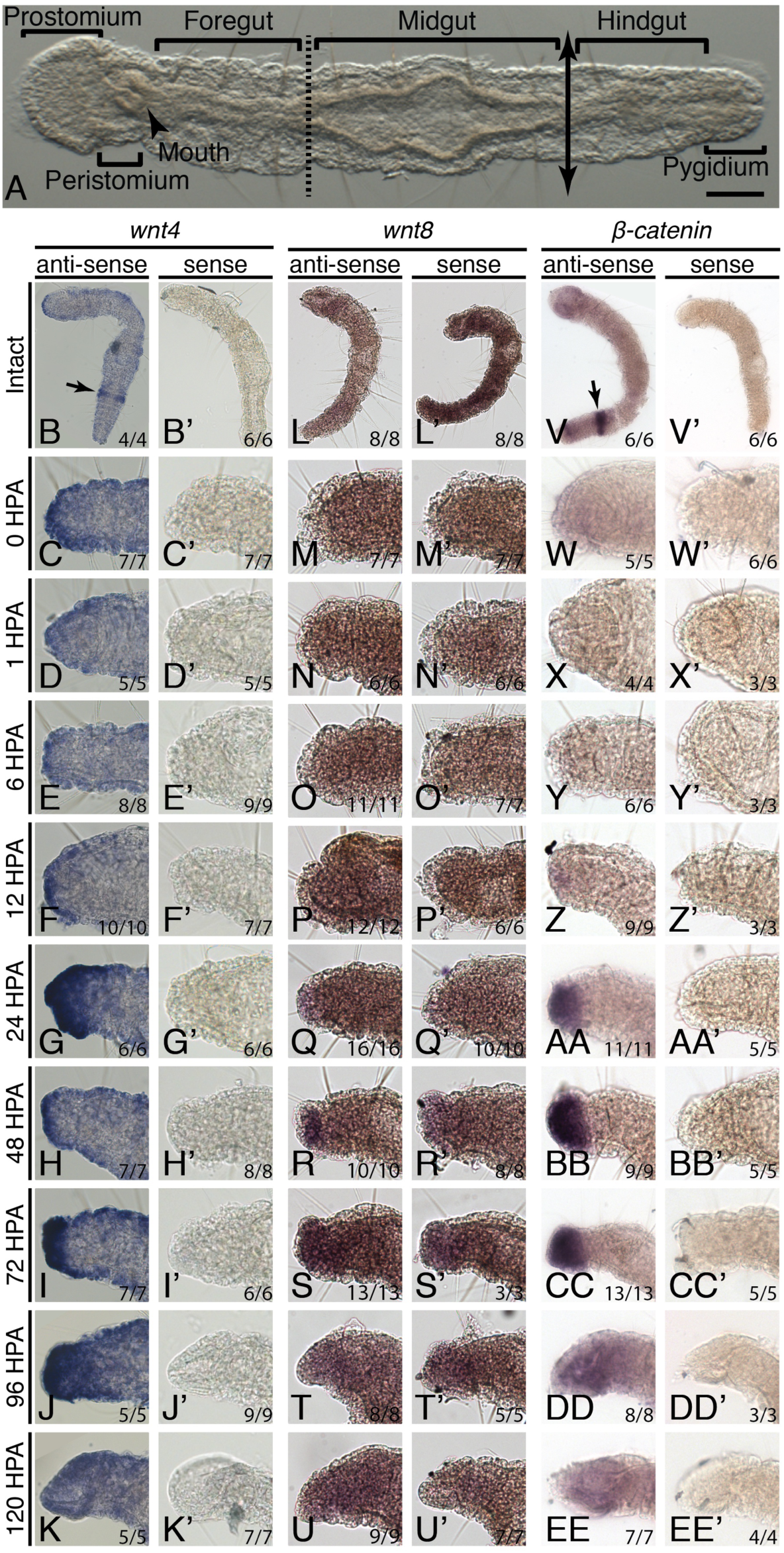
Regenerating wounds show *wnt4* and *β-catenin* activation during blastemal cell proliferation. (**A**) Dorsal view of an intact worm. Double arrow line indicates posterior synchronizing cutting plane between midgut and hindgut junction. Dotted line indicates the anterior amputation plane between foregut and midgut junction. (**C**-**K’**) *wnt4* expression pattern during regeneration. Epithelial *wnt4* signal decreased within the first 6 HPA (**C**-**E**), followed by activation in the blastema between 24-96 HPA (**G**-**J**). (**M**-**U’**) *wnt8* expression in the blastema was indistinguishable from background signals. (**W**-**EE’**) *β-catenin* expression pattern during regeneration. *β-catenin* was activated in the blastema between 24-72 HPA (**AA**-**CC**). In intact worms, *wnt4* and *β-catenin* could be detected at the posterior proliferating segments of intact worms (arrows of **B** and **V**). Ratio at the lower right corner indicates specimens showing the corresponding expression patterns. Scale bar in **A** = 100 μm.

In this report, we first showed transcriptional activation of *wnt4* and *β-catenin* during regeneration. Then, to assay Wnt pathway activity, we labeled nuclear localized β-catenin protein by immunohistochemistry. We found blastemal formation was accompanied with increasing levels of nuclear β-catenin between 12-24 HPA, which was followed by dropping levels of nuclear β-catenin when anterior structures gradually reformed. To functionally interrogate the pathway, we treated the worms with chemical inhibitors and found that both activating and inhibiting the pathway impaired regeneration. Further short-term treatments demonstrated that the chemical activator blocked wound closure and blastema formation; while the inhibitor affected neuron and muscle regeneration in the blastema. Lastly, we labeled proliferating cells and found that cell proliferation in wound tissue was inhibited when over-activating Wnt pathway. The dynamic changes of nuclear β-catenin level and discrete cellular responses under inhibitor treatments suggest that wound closure, blastemal formation and cell differentiation depends on mild activation of endogenous Wnt pathway. Tilting the balanced pathway activity to either extremity impaired complete tissue regeneration.

## Results

### Wnt pathway is activated in the regenerating blastema

To investigate whether Wnt pathway is involved in regeneration, we first tested the spatiotemporal expression patterns of *wnt4, wnt8* and *β-catenin* by whole-mount *in situ* hybridization (WMISH, Fig. 1). Upon amputation, the cutting plane was immediately covered by *wnt4* expressing epithelial cells (Fig. 1C). Then the epithelial *wnt4* signal diminished from the first hour-post-amputation (HPA) to 6 HPA before the gene was activated again at the wound from 12 HPA (Fig. 1D-F). The *wnt4* signal persisted in blastema until head structures completely reformed at 120 HPA (Fig. 1K). On the contrary, we did not detect significant up-regulation of *wnt8* throughout the repairing process (Fig. 1M-U’), suggesting preferential Wnt ligand activation during regeneration. *β-catenin* was activated from 24 HPA through 72 HPA and diminished when nascent head structures became recognizable at 96 HPA (Fig. 1DD).

Additionally, we observed *wnt4* and *β-catenin* expression at the posterior 1/4^th^ segment of intact worms (Fig. 1B and V) where asexual fission occurs and EdU labeled proliferating cells are enriched (C.-P. Chen et al., 2020; Falconi et al., 2015) The expression of *wnt4* and *β-catenin* at this domain suggest that these genes may regulate the development of posterior growth zone, like other multipotent genes, such as *piwi*, *vasa* and *nanos* in other annelids (Gazave et al., 2013; Kozin & Kostyuchenko, 2015; Zheng et al., 2016).

In most studied models, active Wnt signaling cascade stabilizes cytoplasmic pool of β-catenin protein, which then translocate and accumulate in the nuclei and mediate downstream gene transcription (Clevers & Nusse, 2012). We labeled β-catenin protein by immunohistochemistry to evaluate the pathway activity during regeneration (Fig. 2). Combining with nuclear staining, we found β-catenin localized in the nuclei of regenerating blastema (Fig. 2A-D). Because nascent blastema was hardly distinguishable during early wound closure, we quantified the ratio of β-catenin-enriched (β-catenin^+^) nuclei to the blastemal nuclei from 12 HPA (Fig. 2R). We found that when the proliferating blastema gradually became distinguishable from wound epithelium between 12 to 24 HPA, the ratio of β-catenin^+^ nuclei increased. Then the ratio of β-catenin^+^ nuclei decreased as head regeneration completed. Combined with ISH data, we proposed that Wnt signaling is activated in the proliferating blastema from 12 HPA and is gradually silenced until regeneration completes at 120 HPA.

**Fig. 2.**
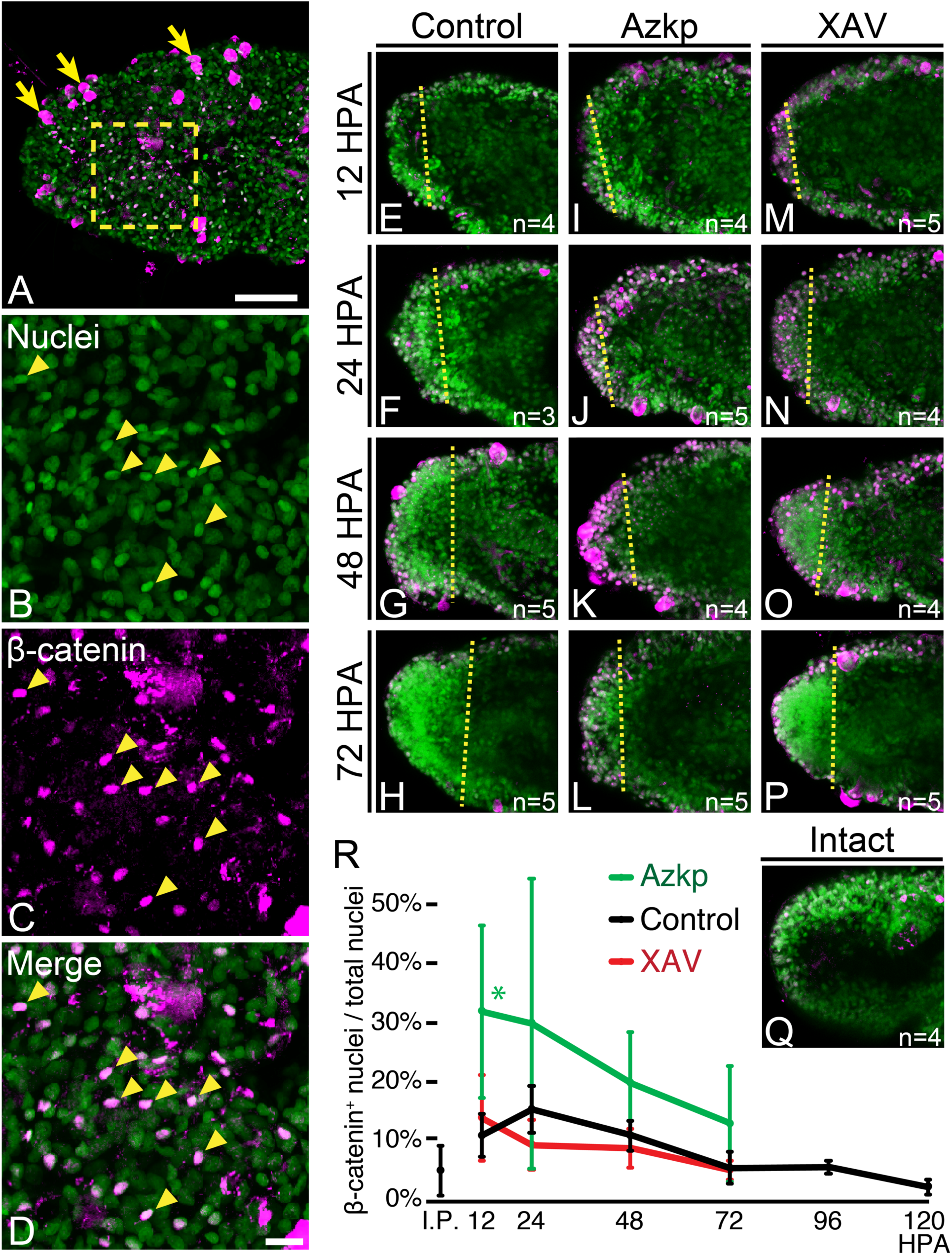
Nuclear β-catenin suggests Wnt pathway activation during blastemal proliferation at 24 HPA. (**A**-**D**) Ventral view of 24 HPA blastema shows β-catenin protein (*magenta*) accumulated in some nuclei (*green*) as indicated by arrowheads. Some non-specific signals were also detected on the epithelium (arrows). (**B**-**D**) Higher magnification of the box region in **A**. (**E**-**P**) Representative images of nuclear β-catenin at regeneration stages under control or inhibitor treatments. (**Q**) Intact prostomium of β-catenin immunohistochemistry stain. (**R**) The ratio of β-catenin^+^ nuclei to total blastemal nuclei under different conditions. Control blastema showed highest level of β-catenin^+^ at 24 HPA. Azkp enhanced nuclear β-catenin level (*green* line), while XAV down-regulated nuclear β-catenin level at 24 and 48 HPA (*red* line). **E**-**Q** are maximum projections from ten serial focal planes. Error bars represent the standard deviation of the mean from >3 worms. I.P. = Intact prostomium. Scale bar of **C** = 100 μm; **C’** = 10 μm. *Yellow* dotted lines of **E**-**P** indicate the amputation plane. * denotes *p*<0.05 of *t*-test compared with control at the corresponding time point.

### Perturbing Wnt pathway inhibited *A. viride* regeneration

We next asked whether tuning Wnt pathway can modulate *A. viride* regeneration or not. To further test the role of Wnt signaling in *A. viride* regeneration, we incubated amputated worms in chemical activators: Wnt Agonist, Alsterpaullone (a GSK-3β inhibitor) and 1-azakenpaullone (Azkp, a GSK-3β inhibitor) or inhibitor: XAV939 (XAV, a tankyrase inhibitor and, in turn stabilizes axin Broun et al., 2005; Huang et al., 2009; Kunick, Lauenroth, Leost, Meijer, & Lemcke, 2004). Since endogenous Wnt pathway was activated during blastemal initiation, we expected chemically activating Wnt pathway may enhance blastemal proliferation. However, to our surprise, these compounds inhibited *A. viride* regeneration in dose-dependent manners (Fig. 3A-D). For consistency, we applied 0.25 µM Azkp or 5 µM XAV in the following experiments. We found smaller blastema of Azkp treated worms, suggesting over-activating Wnt pathway may inhibit blastema formation (Fig. 3H-J). On the other hand, XAV did not affect blastema formation (Fig. 3K-M), but worms did not regenerate prostomium (Fig. 3G, M). Additionally, we validated the inhibitor effects by assaying the nuclear β-catenin level in Azkp or XAV treated blastema (Fig. 2I-R). As expected, Azkp resulted in more nuclear localized β-catenin, whereas XAV treated blastema showed less nuclear β-catenin than control at 24 and 48 HPA, demonstrating the efficacy of these inhibitors.

**Fig. 3.**
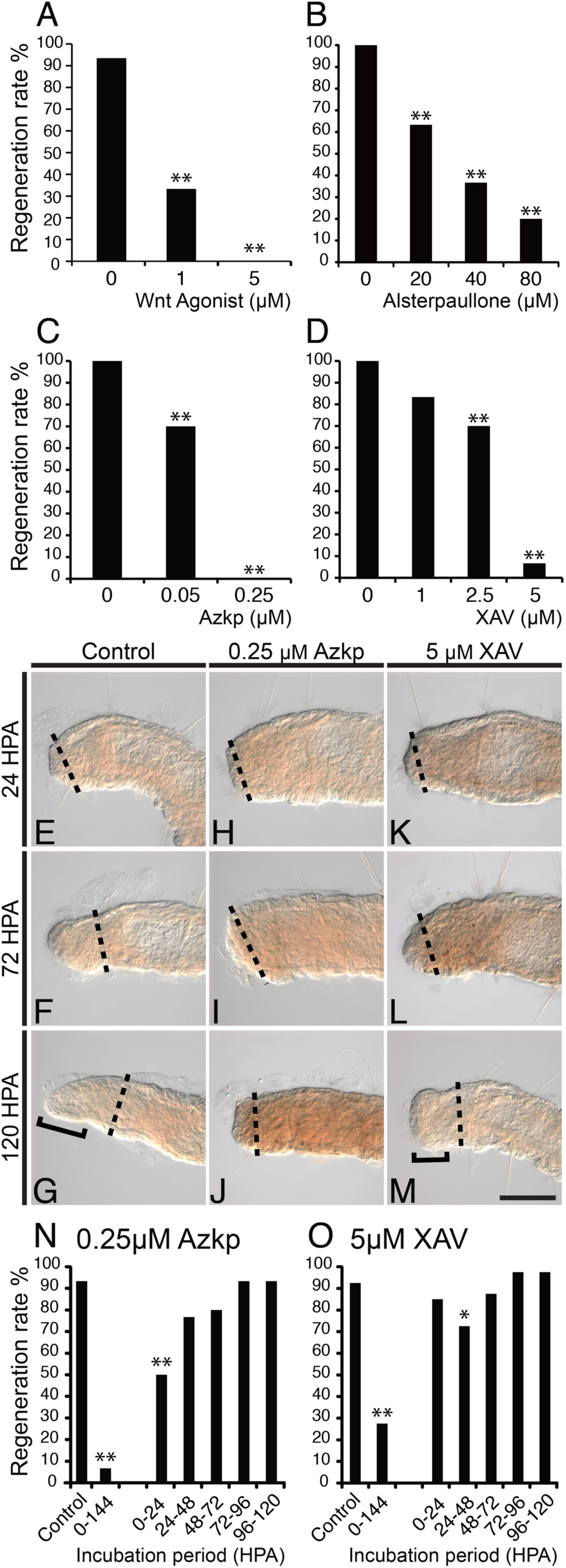
Over-activating and down-regulating Wnt pathway both inhibited regeneration. (**A**-**D**) Exogenous Wnt pathway activators (Wnt agonist, Alsterpaullone and Azakenpaullone), and inhibitor (XAV939), inhibited regeneration in dose-dependent manners. (**E**-**M**) Lateral view of regeneration stages under control, 0.25 μM Azkp or 5 μM XAV treatments. Azkp treated worms showed smaller blastema (**H**-**J**). XAV939 did not affect blastema formation (**K**-**M**), but worms did not regenerate prostomium (compare bracket regions of **G** and **M**). (**N**-**O**) Successful regeneration rates under 24H pulses. Regeneration was significantly inhibited by 0.25 μM Azkp treatment between 0-24 HPA (**N**), and by 5 μM XAV treatment between 24-48 HPA XAV (**O**). Anterior end is toward the left and dorsal to the top. Dotted lines indicate cutting planes. Scale bar in **M** = 100 μm. All images are at the same scale. * denotes *p*<0.05. ** denotes *p*<0.01.

In the above assays, we applied treatments throughout regeneration and found both Wnt pathway activator and inhibitor impaired regeneration. From nuclear β-catenin level of control worms, we found dynamic Wnt activities during regeneration. We hypothesized that different Wnt pathway activities may be required at different regeneration stages. To test the hypothesis, we applied 24H pulse of Azkp or XAV for detailed temporal examinations (Fig. 3N-O). We found treatments of Azkp and XAV before 72 HPA resulted in greater incidences of defective regeneration, suggesting the activity of Wnt pathway is required during early regeneration stages. However, the two treatments showed different effective time windows: 0-24 HPA amputees were sensitive to Azkp, while 24-48 HPA amputees were sensitive to XAV. Combining nuclear β-catenin data with 24H pulse treatments, we propose that wound tissue may require low Wnt pathway activities during initial 0-24 HPA and then high activities between 24-48 HPA.

### Perturbing Wnt pathway impaired neuron and muscle regeneration

Our key criteria for assaying anterior regeneration of *A. viride* are the reformation of head structures (the prostomium, the horseshoe-shaped mouth in the peristomium, Fig. 1) and free gliding behavior on substrates. Since the majority of Azkp and XAV treated amputees failed to meet these criteria, we asked whether the neuron or muscle regeneration is impaired. We first examined the underlying innervation by immunolabeling acetylated α-tubulin (Fig. 4). After five days of regeneration, control *A*. *viride* recovered anterior neuron connections between the neuropils and the ventral nerve cords through the circumesophageal commissures, as well as dorsal-anterior sensory organs of the prostomium (Fig. 4A, D; Hessling & Purschke, 2000). Azkp treated worms failed to regenerate prostomium, and the ventral nerve cords of the remaining tissue showed degeneration and did not extend anteriorly to form circumesophageal commissures or neuropils (Fig. 4E, H). The XAV treated worms showed partial recovery of the neuron connection from the ventral nerve cords to the neuropil through the circumesophageal commissures (Fig. 4I, L). However, the prostomium and the innervation from neuropil to the dorsal-anterior sensory organs were missing, which may be the main reason for the failure of behavioral recovery.

**Fig. 4.**
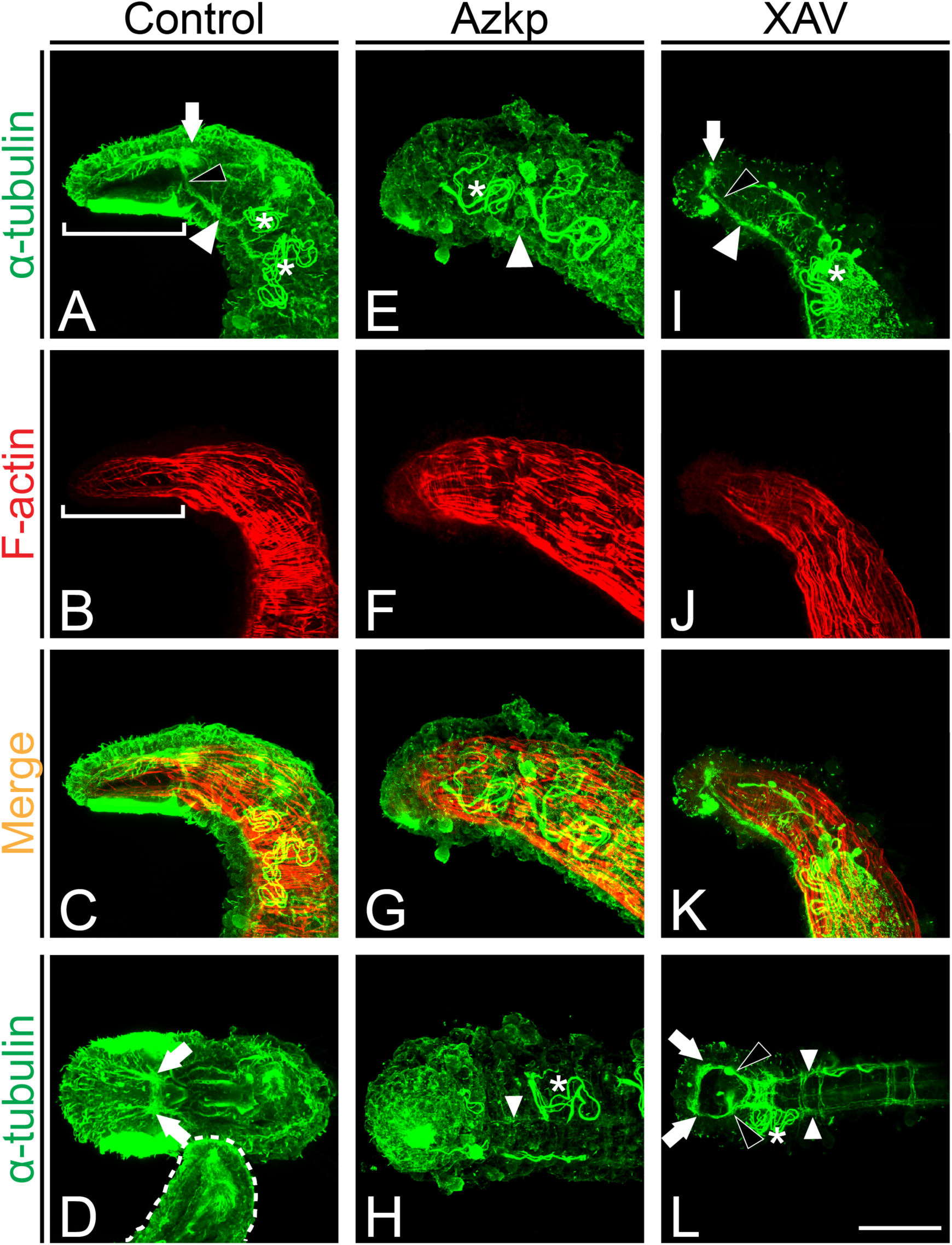
Over-activating Wnt pathway inhibited neuron regeneration and maintenance. (**A**-**D**) By 144 HPA, control worms showed recovered innervation (acetylated α-tubulin, *green*) with neuropil in the prostomium (*white* arrows) connecting to the ventral nerve cord (*white* arrowhead) by the circumesophageal commissures (*black* arrowhead). The neuropil extended exons dorsal-anteriorly to the epithelial sensory cilia. Ventral prostomial ciliary field was also enriched with acetylated α-tubulin (bracket of **A**). Rhomboid shape-arrayed muscle net (F-actin, *red*) supported the flat prostomium (bracket region of **B**). The *white* dotted line in **D** delineates the pygidium of the worm. (**E**-**H**) Azkp treated worms did not regenerate anterior innervations from the degenerated ventral nerve cord (*white* arrowheads) and the prostomium (**F**). (**I**-**L**) XAV treated worms regenerated anterior innervation where the neuropil (*white* arrows) and circumesophageal commissures (*black* arrowheads) connect to the ventral nerve cords (*white* arrowheads). However, these worms did not regenerate the prostomium (**J**). Asterisks indicate metanephridia. Anterior of all amputees are toward the left. Figure **A**-**C**, **E**-**G**, and **I**-**K** are lateral views with dorsal to the top. Figure **D** is dorsal view; **H** and **L** are ventral views. Scale bar in **L** =100 μm. All images are at the same scale.

The rhomboid shape-arrayed muscle net underlying the intact prostomium supports its flat shape and multi-directional movements (Fig. 4B). Such specialized musculature of prostomium can be distinguished from the rest of body segments, which is composed by the perpendicular-arrayed longitudinal and circular muscles. In addition to the observed neuronal regeneration defects, both Azkp- and XAV-treated worms did not regenerate the characteristic prostomium musculature (Fig. 4F, J), which further supported the observed regeneration defects from morphology and behavior.

### Over-activating Wnt signaling by Azkp resulted in blastemal cell proliferation defects

From our previous report, we observed that wound blastema increased in size and showed the most cell proliferation between 24 and 48 HPA (Fig. 3E-G; C.-P. Chen et al., 2020). At the same stages, Azkp treated wound blastema did not show comparable size change like control or XAV treated worms (Fig. 3E-M). Despite Azkp and XAV treatments both inhibited anterior regeneration, subtle differential cellular responses to Wnt activity perturbation may exist. Because the blastema of Azkp treated wounds were smaller, we hypothesized that cell proliferation can be impaired in these worms. We first examined cumulative cell proliferation by 5-ethynyl-2ʹ-deoxyuridine (EdU) pulse between 24 and 48 HPA (Fig. 5B, G, L). EdU^+^ cells at the blastema were less condensed in Azkp treated worms than in control and XAV, suggesting that over-activating Wnt compromised blastemal cell proliferation and resulted in smaller blastema. Additionally, we observed that the EdU^+^ cells were ectopically enriched in the 2^nd^ segment of Azkp treated worms (Fig. 5G). From single z section images, we found the secondary EdU^+^ cell domain located at the anterior tip of the midgut (Fig. 5J), which was observed to lesser extents in control and XAV treated worms (Fig. 5E, O-P).

**Fig. 5.**
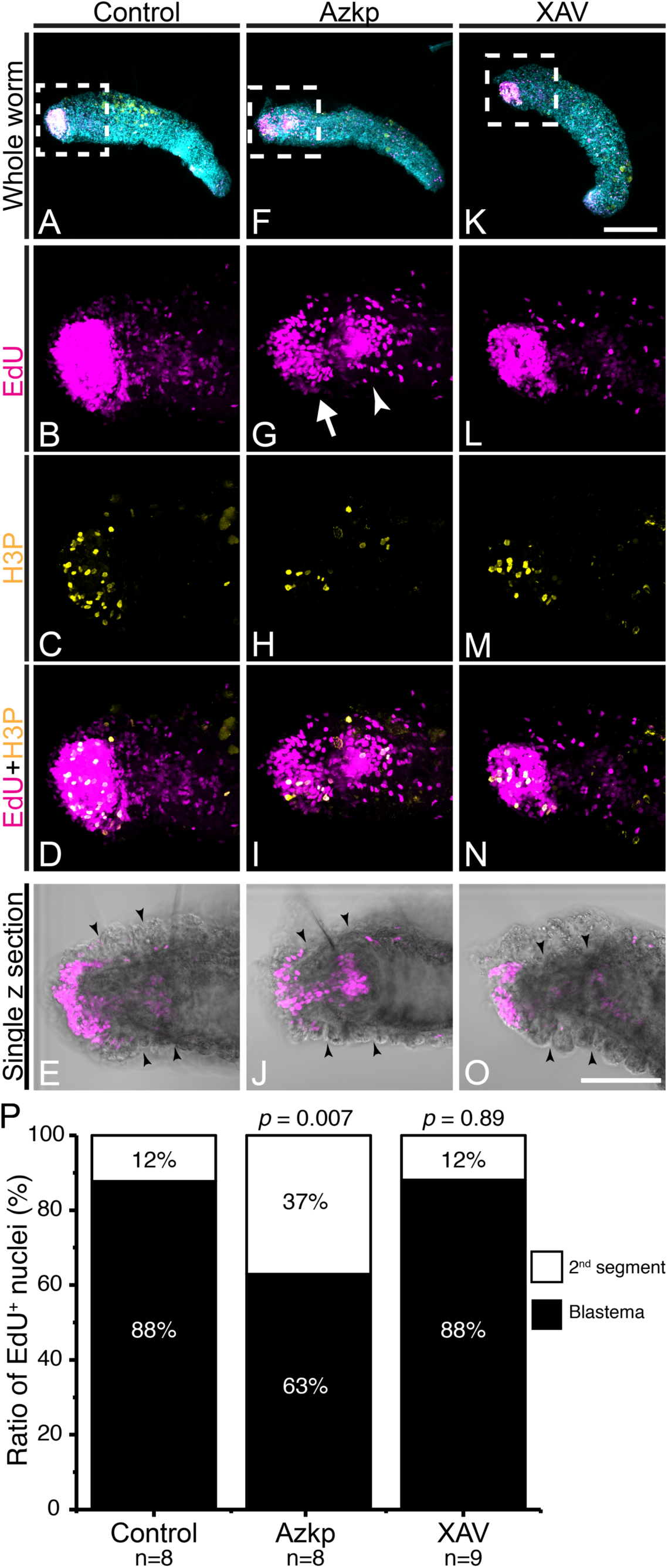
Over-activating Wnt pathway impaired blastemal cell proliferation. (**A**-**O**) Proliferating cells between 24 to 48 HPA are labeled by EdU (*magenta*) or anti-H3P antibody (*yellow*). The nuclei are false-colored in *cyan*. Higher magnification views of *white* rectangle areas of **A**, **F** and **K** are in the corresponding lower panels. In control and XAV treated worms, active proliferating blastema showed condensed EdU signals (**B** and **L**); while Azkp treatment resulted in less EdU^+^ nuclei at the wound site (arrow) and a second domain of proliferating cells in the 2^nd^ segment (arrowhead, **G**). (**J**) Single z section image shows that the proliferating cells in the 2^nd^ segment were located at the front end of midgut. Anterior of all amputees are toward the left with dorsal to the top. Black arrowheads in **E**, **J** and **O** delineate the blastema and the segment boundary. (**P**) Ratio of EdU^+^ nuclei between blastema and the following 2^nd^ segment. Student *t-*test comparison with control worms shows significant (*p* = 0.007) enrichment of EdU^+^ nuclei at the 2^nd^ segment of Azkp treated blastema. Scale bar in **K** = 200 μm; **O** = 100 μm. **A**, **F** and **K** are at the same scale. **B-E**, **G-J** and **L-O** are at the same scale.

Additionally, we also labeled proliferating cells at 48 HPA by anti-phospho-histone 3 antibody (H3P, Fig. 5C, H, M). By quantifying the H3P^+^ cells, we found the number of proliferating cells was significantly lower in the Azkp treated blastema than control and XAV treatments (Control: 55±33.4; Azkp: 18.8±8.2; XAV: 59.2±30.7; *p* of Control-Azkp = 0.01; *p* of Control-XAV = 0.89; *p* of Azkp-XAV = 0.001; Mann-Whitney U-test). Combining EdU and H3P assays, Azkp-induced blastemal formation defects can be attributed to lacking proliferation at blastema and mis-regulation of proliferating domain at the 2^nd^ segment.

## Discussion

In this report, we interrogated the involvement of Wnt pathway in *A. viride* anterior regeneration. Wnt pathway activities were dynamic during regeneration, with the highest activity at 24 HPA when the wound blastema was actively proliferating. Long term incubation with either chemical inhibitor or activator of Wnt pathway impaired regeneration. Further temporal experiments suggested the requirements for the relatively low Wnt activity between 0-24 HPA and high activity between 24-48 HPA. Examinations of innervation, musculature and cell proliferation revealed the underlying cellular responses under treatments. By integrating current observations, we provide the following reasonings about how Wnt pathway is involved in *A. viride* regeneration.

### *A. viride* regeneration depends on dynamic Wnt signaling activities

The dynamic nuclear β-catenin levels suggest Wnt pathway activity varies through stages of *A. viride* regeneration. Thus, long-term treatments by either Azkp or XAV may counteract against endogenous Wnt activities at different stages and result in the ultimate regeneration failure. Then, we further narrowed the treatment into 24H pulses. We found that the two chemical inhibitors were effective before 72 HPA when normal wound tissue repaired through wound closure, blastemal formation and partial differentiation stages. Immediately after amputation, we observed that the wound was covered by nearby *wnt4* expressing epithelium, and then *wnt4* expression was down-regulated from 1 HPA until being activated again at the blastema from 12 HPA. Downregulation of *wnt4* at the wound site suggests that the wnt4-mediated Wnt signaling was silenced during wound healing. Therefore, Azkp treatment at this stage may counteract the Wnt inactivation as well as compromise the following blastemal cell proliferations. Alternatively, over-activating Wnt pathway may override endogenous activation between 12-24 HPA and turn the tissue into non-permissive conditions for regeneration. Then, XAV inhibited regeneration during 24-48 HPA when the most β-catenin accumulated in the nuclei. Although blastemal proliferation was not affected by XAV, the underlying innervation and prostomium musculature could not completely recover. Because the appearance of neuron and muscle cells are signatures of blastemal cell differentiation from nascent stem cells, we propose that the endogenous Wnt activation at 24 HPA can be required for blastemal cell differentiation.

Collectively, as summarized in Fig. 6, our data suggest that the progression of regeneration is dependent on different Wnt activities: wound closure and initial repairing occur under low Wnt activity (0-24 HPA); high Wnt activity initiates blastemal proliferation (24 HPA); blastemal differentiation requires median Wnt activity (24-48 HPA); continued completion of regeneration after 72 HPA is inert to exogenous Wnt activity modulators.

**Fig. 6.**
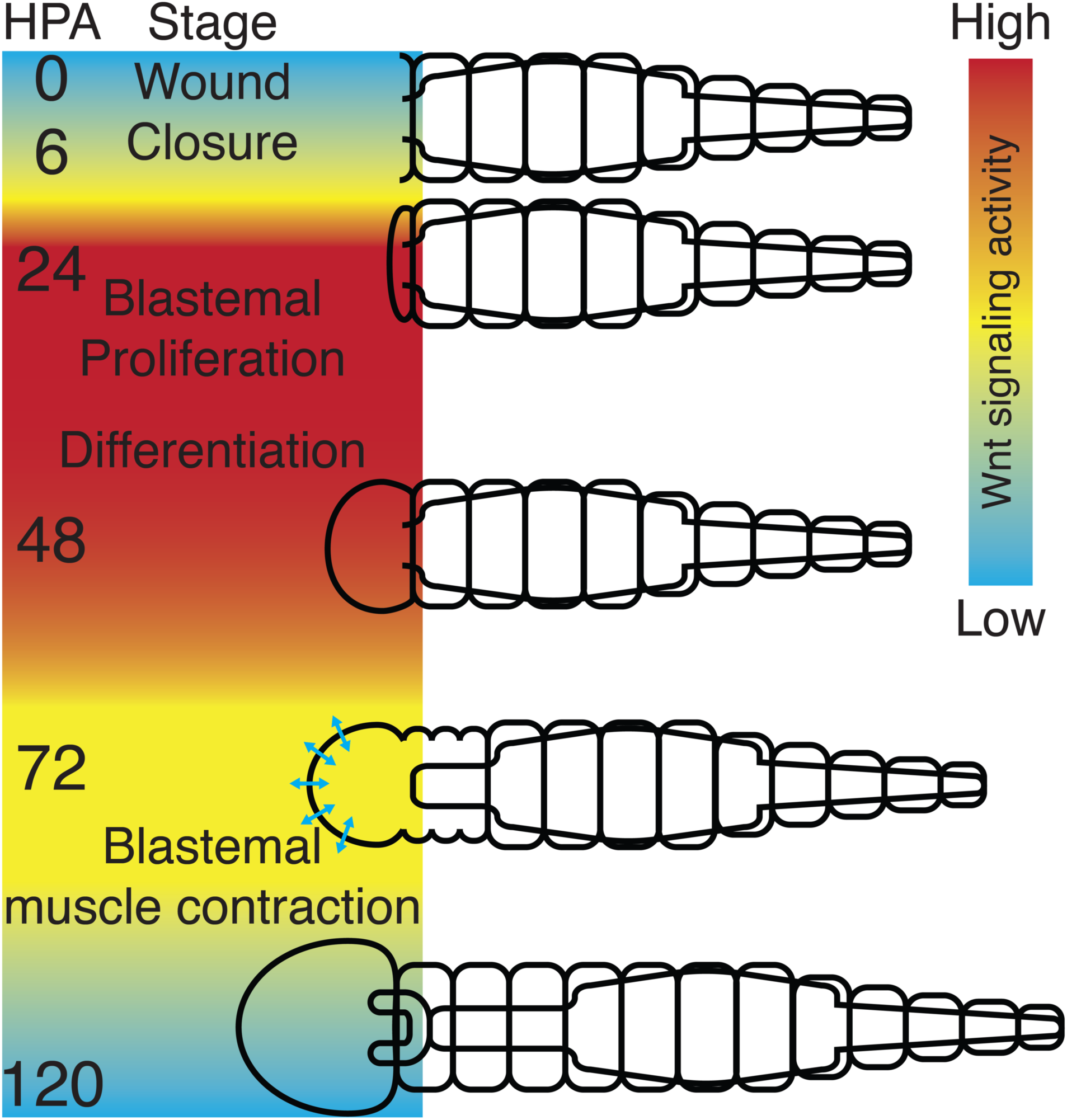
A proposed model of endogenous Wnt pathway activity during *A. viride* regeneration. A rainbow pallet defines hypothesized blastemal Wnt activities. After amputation, wound closure finishes in the first 6 HPA. Then blastema gradually enlarges at the wound site by cell proliferation. By 72 HPA, blastemal tissue initiates occasional contraction, indicating neuron and muscle cell differentiation. By 120 HPA, worms complete regeneration with a horseshoe-shaped mouth and can freely glide on substrates.

### Wnt signaling may not regulate anterior-posterior identity of regenerating *A. viride*

Wnt pathway is indispensable for defining axis during embryogenesis and regeneration of many animal models (Broun et al., 2005; Chera et al., 2009; Gurley et al., 2008; Iglesias et al., 2008; Leclère, Bause, Sinigaglia, Steger, & Rentzsch, 2016; Lengfeld et al., 2009; Petersen & Reddien, 2008; E. Tanaka & Weidinger, 2008). In planarian, activating Wnt pathway promotes posterior regeneration at the anterior wound; while inhibiting the pathway induces head formation at the posterior wound (Iglesias et al., 2008; Petersen & Reddien, 2008). If axial patterning of Wnt pathway is evolutionarily conserved in *A. viride*, we anticipated that anterior regeneration would be enhanced when we inhibited the pathway, and that posterior regeneration would be enhanced when we over-activated the pathway. However, our current data and methods do not support such hypothesis, i.e. XAV did not enhance anterior regeneration. Additionally, we did not observe ectopic head regeneration from the posterior wounds from our preliminary XAV treatments. Prostomium and anterior neuronal regeneration were inhibited by XAV treatments, suggesting other side effects existed. On the other hand, anterior regeneration was inhibited when the pathway was over-activated by Azkp, but we could not confidently claim that these Azkp treated wounds were transformed to posterior identity for lacking axial markers. Therefore, labeling anterior-posterior molecular markers will be the future direction for testing whether Wnt pathway regulates the axial identity of regenerating *A. viride*.

### GSK-3β activity is required for *A. viride* neuronal regeneration and maintenance

Innervation to the regenerating tissues can be necessary and sufficient for complete regeneration in many systems (A. Kumar & Brockes, 2012). For example, bisected salamander limbs cannot regenerate if the remaining neuron is removed (A. Kumar & Brockes, 2012); ectopic neuron growth initiates ectopic blastemal formation in annelids (Bely, 2014). In planarian, β-catenin-1 localizes in the nuclei of brain and ventral nerve cords and is required for brain patterning during regeneration (Hill & Petersen, 2015; Sureda-Gómez, Martín-Durán, & Adell, 2016). Azkp treatment results in neuronal regeneration defects in planarian, such as smaller cephalic ganglia and ectopically projected visual axons (Adell, Marsal, & Saló, 2008). Consistent with this idea, the Azkp treated *A. viride* did not regenerate neurons and showed degenerating ventral nerve cords, suggesting the requirement of GSK-3β in neuronal regeneration and maintenance. On top of that, planarian neuron regeneration is required for blastemal morphogenesis (Adell et al., 2008). In *A. viride*, the blastemal cell proliferation and innervation were both impaired when GSK-3β was inhibited by Azkp, suggesting a linkage between neuron regeneration and blastema formation. If neuron is generally required for regeneration in diverse animal models, then it is likely that *A. viride* blastema formation can be also regulated by innervation, which possibly requires active GSK-3β. However, the epistatic or mutual regulation of innervation and blastema formation in *A. viride* is not clear yet and warrants further investigations.

In summary, we demonstrated that proper regulation of Wnt signaling activities is required for different stages of *A. viride* anterior regeneration. As an echo to other regeneration models, our data further generalize the involvement of Wnt signaling to regeneration in an evolutionarily conserved perspective. In the future, more sophisticate approaches, such as CRISPR/Cas9 mutagenesis, will provide detailed molecular mechanisms of annelid regeneration as well as comparative regeneration biology in a broader context.

## Materials and methods

### Animal husbandry

Animal husbandry followed Chen et al. 2020. The whole culturing colony was asexually descended from one single worm to avoid genetic differences.

### Amputation

Before anterior amputation, *A. viride* was first bisected between the midgut and hindgut junction (Fig. 1A). By this method, we aimed to synchronize the posterior segments and to prevent any proliferating progeny interfering with assessing successful regeneration (C.-F. Chen, Sung, Chen, & Chen, 2018). During synchronization, *A. viride* was cultured in ASW at 25°C without feeding. The pygidium recovered in three days. Then, the anterior segments were amputated between the foregut and midgut junction (Fig. 1A).

By 144 hour-post-amputation (HPA), successful anterior regeneration was assessed by the following criteria: **1**) muscle contraction in prostomium and peristomium, **2**) appearance of horseshoe-shaped mouth (Fig. 1A); **3**) voluntary gliding behavior on substrates. Regeneration rates were assayed by averaging the proportion of successful regeneration worms form at least 3 biological replicates. Each replicate was started from >10 worms at 0 HPA.

### Isolation of *A. viride wnt4*, *wnt8*, and *β-catenin* sequences

Total RNA of *A. viride* was purified from intact worms by TRIzol (Thermo Fisher Scientific; Waltham, MA) following manufacturer’s instructions. Then, the cDNA library was reverse-transcribed by SuperScriptIII Reverse Transcriptase (Thermo Fisher Scientific) with oligo dT as the first strand primer. Using cDNA library as template and degenerated primers (Table 1), we amplified target fragments by PCR, cloned the fragments into T&A^TM^ vector (Yeastern Biotech Co., Taiwan), and sequenced the insert fragment.

**Table 1.**
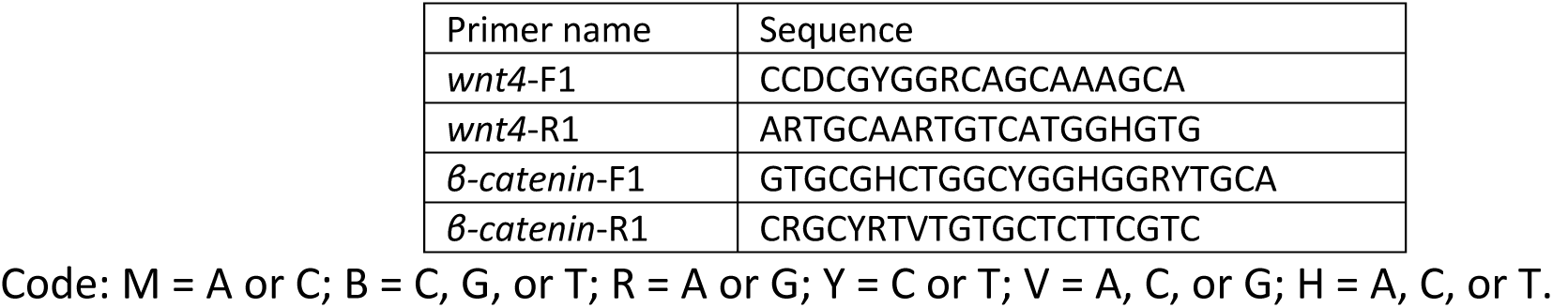
Deg nerate primers initially used to clone Wnt pathway related genes.

For 3’ Rapid Amplification of cDNA Ends (3’ RACE), the cDNA library was primed by 3’ end anchor primer (3’ AP, Table 2). Then gene specific internal primers and 3’ AP head primer were used for nested PCR (Table S2). The amplified products were cloned into T&A^TM^ vector and sequenced.

To amplify the 5’ end sequence, we performed 5’ RACE by reverse transcribing the first strand cDNA by gene specific primer (Table 2). Then multiple deoxycytidines were added to the 3’ end of cDNA by terminal deoxynucleotidyl transferase (ThermoFisher). The gene specific primers and universal 5’ end anchor primer (5’ AP, Table 2) were used to amplify the 5’ end sequence. Secondary nested PCR with 5’ AP head primer (Table 2) was used to further amplify the specific sequence, which is followed by cloning and sequencing as described above.

**Table 2.**
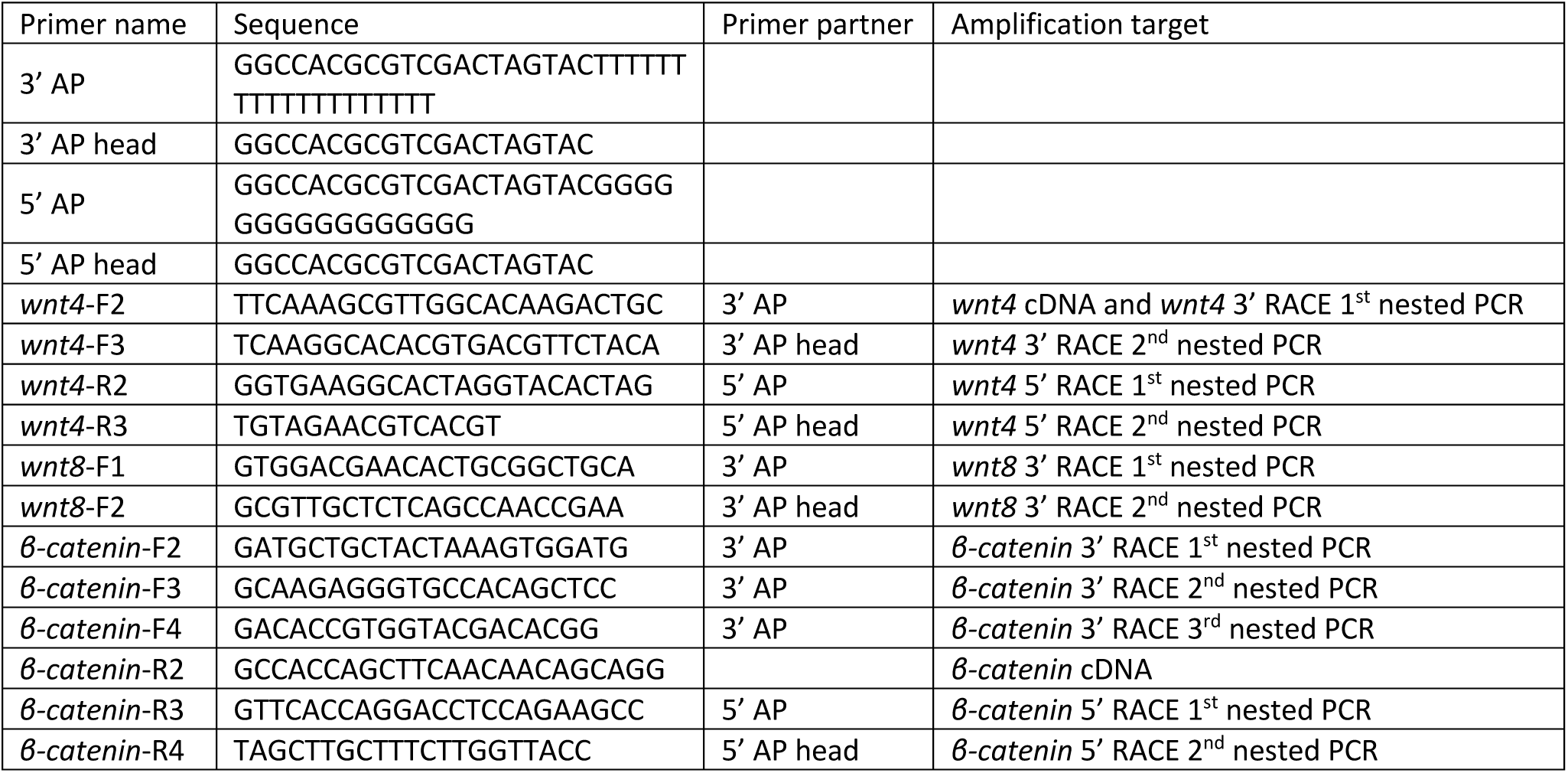
Primers for 5’ and 3’ RACE. Note that *wnt4*-R2 and *β-catenin*-R2 were used as gene specific primers for first strand cDNA synthesis during reverse transcription of 5’ RACE.

### Gene homology analysis

Thirteen subfamilies of Wnt homologues proteins were previously identified in Lophotrochozoan genomes (Cho, Vallès, Giani, Seaver, & Weisblat, 2010; Prud’homme, Lartillot, Balavoine, Adoutte, & Vervoort, 2002). To test the homology of *A. viride wnt* genes, we first used SMART protein analysis (http://smart.embl.de/) to identify the WNT domain of Lophotrochozaon Wnt genes and aligned them by the default parameter of MUSCLE of MEGA7 program (S. Kumar, Stecher, & Tamura, 2016). Then, the phylogenetic relationship of wnts were resolved by Neighbor-joining method by p-distance model with 10,000 bootstrap repeats clustering *A. viride wnt4* and *wnt8* within the corresponding subfamilies (Fig. 7).

**Fig. 7.**
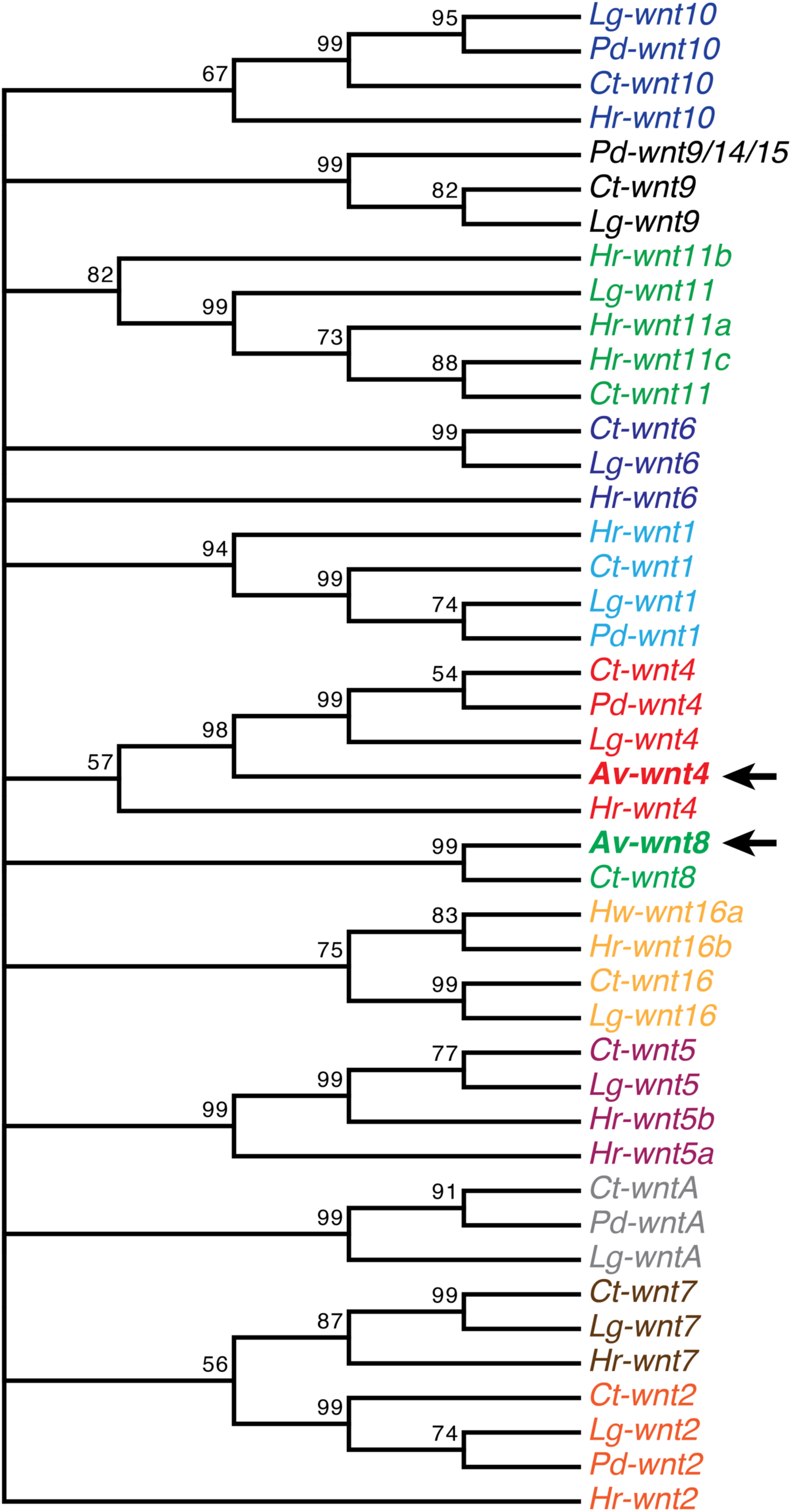
Neighbor-joining tree clusters *A. viride wnt* genes into the corresponding Lophotrochozoan *wnt* subfamilies. *Av-wnt4* and *Av-wnt8* are indicated by arrows. Within the 10,000 replicates bootstrap tests, the percentage of replicate trees in which the associated taxa clustered together are shown at the branches. Note that the nodes with bootstrap percentage <50% are compressed. Species name abbreviations: *Av: Aeolosoma viride*; *Ct*: *Capitella teleta*; *Hr*: *Helobdella robusta*; *Lg*: *Lottia gigantea*; *Pd*: *Platynereis dumerilii*.

The cloned β-catenin protein encodes 12 characteristic armadillo domains (ARM) by SMART analysis. The amino acid sequence of β-catenin showed higher similarity to *β-catenin* than *α-catenin* or *γ-catenin* homologs by NCBI protein BLAST, confirming the sequence identity as *Av-β-catenin*.

### Chemical inhibitor treatments

Anterior amputated *A. viride* were washed with treatment solutions four times before incubating either through whole regeneration process (Fig. 3A-D) or at 24H intervals (Fig. 3N-O). 1-Azakenpaulone (Azkp; MilliporeSigma, Burlington, MA), XAV939 (XAV, MilliporeSigma), Alsterpaullone (MilliporeSigma), Wnt Agonist (MilliporeSigma) were dissolved in dimethyl sulfoxide (DMSO) as stock solutions, and then diluted in artificial spring water (ASW) as working solutions at designed concentrations. Control solution was prepared with comparable concentrations of DMSO in ASW.

### Whole-mount Immunofluorescence (WMIF)

Worms were anesthetized by menthol saturated ASW, and then fixed with 4% PFA-ASW overnight at 4°C. Then the samples were washed 5 times with phosphate buffered saline (PBS, pH=7.4) containing 0.1% Triton X-100 (PBST). After washing, the fixed worms were rinsed with 100% methanol 3 times at RT and stored at −20°C until proceeded to immunohistochemistry.

To begin immunohistochemistry, the samples were re-hydrated through series of 66% and 33% methanol in PBST washes and additional washes in PBST for 5 times. Then, the samples were treated with 5 μg/mL protease K solution at RT for 5 minutes, followed by incubation in 2 mg/mL glycine for 5 minutes and post-fixation with 4% PFA-PBST at RT for 1 hour. After post-fixation, the samples were washed 3 times with PBST, and immersed in the blocking solution (PBST with 3% bovine serum albumin and 5% Goat serum) for up to 1 hour at RT. Primary antibody incubation: Rabbit-anti-β-catenin antibody (in 1:800 dilution; MilliporeSigma, C2206), Rabbit-anti-H3P antibody (in 1:1000 dilution; MilliporeSigma, 06-570) or Mouse-anti-acetylated α-tubulin (in 1:1000 dilution; MilliporeSigma, T7451) was applied overnight at 4°C with blocking solution as diluent. After 6 times washing with PBST, samples were incubated in fluorophore conjugated Goat anti-Rabbit antibody or Goat anti-Mouse antibody (in 1:400 dilution; Thermo Fisher Scientific) for 2 hours at RT with blocking solution as diluent. F-Actin staining was performed on no methanol treated samples as follows: After 1 hour blocking by 1% BSA-PBST at RT, we incubated samples in BODIPY® FL Phallacidin (1:40 dilution in blocking buffer, Thermo Fisher Scientific) at RT for 1 hour. The nuclei were counter-stained with 1 μg/mL DAPI-PBST for 20 minutes at RT. After 4 times washes by PBST, the samples were immersed and cleared in 80% glycerol-PBS with 0.1% NaN_3_ before mounting on slides.

### Whole-mount *in situ* hybridization (WMISH)

DNA templates of riboprobes were amplified by gene specific primers (Table 3) and cloned into T&A^TM^ vector. After confirming the desired orientations by sequencing, we linearized the plasmids as templates for DIG-labeled riboprobes synthesis by T7 polymerase (Thermo Fisher Scientific) with DIG RNA labeling mix (MilliporeSigma).

**Table 3.**
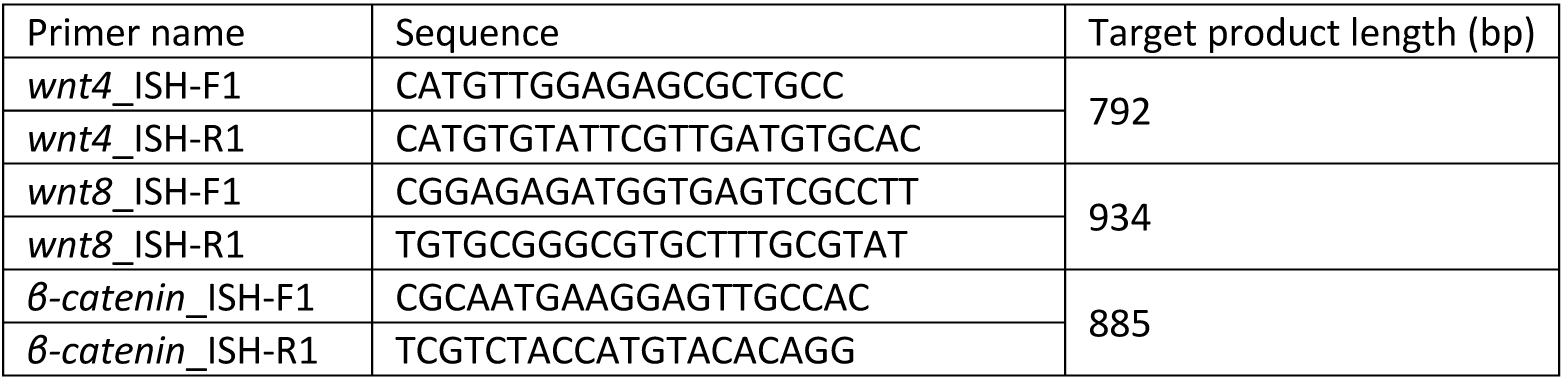
Primers for synthesizing riboprobe templates.

The WMISH protocol was adopted from zebrafish WMISH protocol (Thisse & Thisse, 2008). Sample preparation was similar to WMIF, but PBS with 0.1% Tween-20 (PBSTw) was used instead of PBST. Pre-hybridization was performed in HYB^-^ buffer (50% formamide, 5X SSC, and 0.1% Tween-20 in DEPC-treated H_2_O) at 65 °C for 1 hour. The riboprobes were diluted at 1 ng/µL in HYB^+^ buffer (HYB^-^ buffer with 50 μg/mL Heparin and 0.5 mg/μL yeast tRNA; MilliporeSigma). Hybridization was performed at 65 °C overnight. Then samples were washed at 65 °C by serial dilutions of HYB^-^ and 2X SSCTw (2X SSC with 0.1% Tween-20) from 2:1 to 1:2 ratio, one 2X SSCTw wash, two 0.2X SSCTw (0.2X SSC with 0.1% Tween-20) washes, and another serial dilution of 0.2X SSCTw and PBSTw from 2:1 to 1:2 ratio. The samples were washed once by PBSTw at RT and immersed in blocking solution (5% BSA in PBSTw) at 4 °C overnight. The samples were then incubated in 1:10,000 diluted anti-DIG-AP antibody (MilliporeSigma) in blocking buffer at 4 °C overnight. Then samples were washed 10 times with PBSTw and 3 times with AP buffer (0.1 M Tris-Cl, 0.05 M MgCl_2_, 0.1 M NaCl and 0.1% Tween-20 in DEPC-treated H_2_O). NBT and BCIP were added in AP buffer for colorimetric reaction at 4 °C overnight. Finally, the reaction was stopped by 5 times PBSTw washes and once 100% methanol wash before immersing in 80% glycerol-PBS for mounting.

### EdU labeling

Regenerating *A. viride* were incubated in 100 μM 5-ethynyl-2’-deoxyuridine (EdU; Thermo Fisher Scientific) diluted in ASW from 24 to 48 HPA, when blastemal cells showed the most proliferation (C.-P. Chen et al., 2020). Then, worms were anesthetized and fixed as WMIF method. After protease K treatment and re-fixation, Click-iT® reaction was performed as manufacturer’s instruction (Click-iT® EdU Imaging Kits, Thermo Fisher Scientific). Experiments validated the specificity of EdU is shown in Fig. 8.

**Fig. 8.**
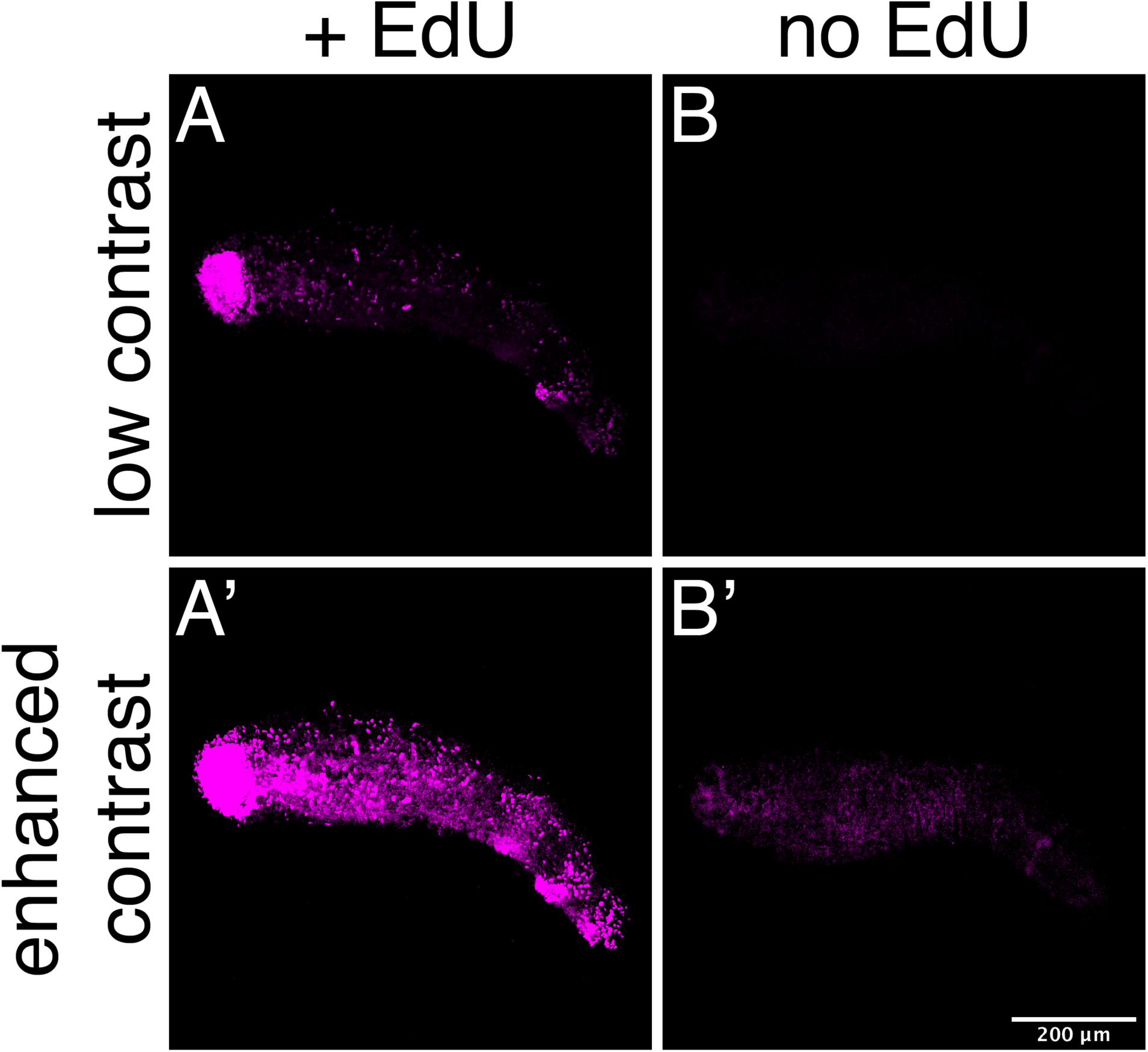
EdU incubation specifically labeled proliferating cells. The regenerating worms were incubated with or without EdU between 24-48 HPA. Without EdU, background signal appeared only when the contrast of image was highly enhanced (**B’**).

### Microscopy

Images of regenerating worms and *in situ* hybridization were taken by Stereo Investigator (MBF Bioscience), Zeiss Axio Observer Z1 or Zeiss Axio Imager A1 microscope with a Zeiss AxioCam MRc CCD camera. Fluorescent specimens were imaged by Leica TCS SP5 confocal laser scanning microscope. Contrasts of images were adjusted as the corresponding control images by ImageJ. Figures were prepared by Illustrator CC (Adobe).

### Image quantification

β-catenin^+^ and H3P^+^ nuclei were manually quantified through z-stacks of regenerating blastema by Fiji: in brief, the images were pre-processed by “Subtract Background”, “Median” and the nuclei were quantified by “3D Maxima Finder” (Boudier et al., 2013). EdU^+^ nuclei quantification was performed according to Cai et. al, 2009 with minor modifications (Cai, Vallis, & Reilly, 2009): analysis of particles was set > 12 µm^2^ corresponding to the cell size and covered the entire z stack range of images.

### Statistic analysis

Difference of successful regeneration rates were tested by Cochran Q test and McNemar change test. The ratio of β-catenin^+^ nuclei was analyzed by two-tailed *t-*test, assuming equal variances. Parametric unpaired *t*-test between treatments was performed for comparing the ratio of EdU^+^ nuclei in blastema to 2^nd^ segment with log-transformation. The number of H3P^+^ nuclei in the blastema was assayed by Mann-Whitney U-test. *p* values less than 0.05 were considered statistically significant.

## Declarations

### Ethics approval and consent to participate

All experimental treatments on *A. viride* were approved by the Environmental Protection and Occupational Safety and Health Center, National Taiwan University.

### Availability of data and material

All data generated or analyzed during this study are included in this published article, or available upon reasonable request from the corresponding author.

### Authors’ contributions

CYC, WTY and JHC prepared the manuscript. CYC and WTY design and perform the experiments and analyzed the data. All authors read and approved the final manuscript.

## Acknowledgements

We would like to thank Dr. Eric Hill from Stowers Institute for Medical Research, Kansas City, MO for suggestions and critics on the manuscript. We appreciate Dr. Jr-Kai Yu from Institute of Cellular and Organismic Biology, Academia Sinica, Taiwan, for kindly providing access to the confocal microscopy.

